# The N terminus-only (*trans*) function of the Adhesion GPCR Latrophilin-1 controls multiple processes in reproduction of *C. elegans*

**DOI:** 10.1101/2023.02.18.529090

**Authors:** Daniel Matúš, Victoria Elisabeth Groß, Franziska Fiedler, Wilbert Berend Post, Alexander Bernd Knierim, Johanna Lena Schön, Torsten Schöneberg, Simone Prömel

**Affiliations:** Rudolf Schönheimer Institute of Biochemistry, Medical Faculty, Leipzig University, Leipzig, Germany; Institute of Cell Biology, Department of Biology, Heinrich Heine University Düsseldorf, Düsseldorf, Germany; Department of Molecular and Cellular Physiology, Stanford University, Stanford CA, USA; Leipzig University Medical Center, IFB Adiposity Diseases, Leipzig, Germany

**Keywords:** Adhesion GPCRs, Latrophilin, *trans* function, germ cells, sperm guidance, apoptosis, proliferation

## Abstract

Adhesion G protein-coupled receptors (aGPCR) are unique molecules. They are able to transmit classical signals via G-protein activation (7TM-dependent/*cis* signaling) as well as to mediate functions solely through their extracellular N termini, completely independent of the seven transmembrane helices domain (7TM) and the C terminus (7TM-independent/N terminus-only/*trans* function). This dual mode of action is highly unusual for GPCRs and allows for a plethora of possible cellular consequences. However, the physiological implications and molecular details of this N terminus-mediated signaling are not well understood. Here, we identify three distinct 7TM-independent/*trans* functions of the aGPCR Latrophilin homolog LAT-1 in the nematode *Caenorhabditis elegans* together regulating reproduction: sperm guidance, germline apoptosis and proliferative activity of germ cells in the gonadal stem cell niche. In these contexts, the receptor elicits its functions in a non-cell autonomous manner from adjacent somatic cells. These functions might be realized through alternative splicing of the receptor specifically generating N terminus-only variants. Thus, our findings shed light on the versatility of 7TM-independent/N terminus-only/*trans* functions of aGPCR and discusses possible molecular details.

## INTRODUCTION

Cellular communication mediated by G protein-coupled receptors (GPCR) typically relies on the transduction of an extracellular cue into a cell, which is mostly realized through the intracellular activation of G proteins via the seven transmembrane helices domain (7TM) of the receptor. This concept has recently also been confirmed for the class of Adhesion GPCR (aGPCR) ^1,2^. Like many other GPCR, these receptors play essential roles in various physiological processes, but harbor structural features that distinguish them as a separate class within the GPCR superfamily (summarized in ^3^). One of these features is their extraordinarily long extracellular N terminus comprising various domains and enabling the receptors to mediate signals and engage in cell-cell or cell-matrix adhesion (summarized in ^3^).

Remarkably and in contrast to other GPCR, members of the class of aGPCR are capable of transmitting functions only via these complex N termini, indicating non-canonical signaling mechanisms completely independent of their 7TM and C terminus. Thereby, these receptors cannot only transduce extracellular cues directly into the cell they are expressed on by classic G protein-mediated cascades requiring the entire receptor molecule (7TM-dependent/ *cis* signaling). They are also able to elicit functions solely via their extracellular N termini that mostly affect adjacent cells (N-terminus-only/7TM-independent/ *trans* function). This dual mode of action is highly uncommon for receptors, especially GPCR. Our work and others’ suggest that several aGPCR, such as Latrophilins/LPHN/ADGRL ^4,5^, BAI1/ADGRB1 ^6^, GPR126/ADGRG6 ^7^, and CD97/ADGRE5 ^8^ harbor the ability to act in such bidirectional manner, and there is evidence for more ^9-11^. These cases imply that the unique dual mode of function is a common feature of this receptor class and data from several studies indicates that the sole N terminus independent of the 7TM and C terminus is always sufficient to mediate effects ^4-8^. However, while the understanding of the basic characteristics of the N terminus-only/7TM-independent function and its cellular consequences are beginning to take shape, essential questions regarding the molecular models of this concept remain unanswered. For instance, while it was observed that this mode of function affects cells surrounding the cells with the aGPCR, it is not shown whether the receptor is activating signaling cascades in these cells, or if its non-cell autonomous effect is mediated – for example – via adhesion. Further, it is unclear whether the whole receptor molecule is required at all times to fulfill both, classical G protein-mediated and N terminus-only functions and whether the N terminus is attached to the entire receptor molecule, released or produced independently of the C-terminal parts to mediate its function. Autocatalytic cleavage at the G protein-coupled receptor proteolysis site (GPS) ^12^ yielding an N-terminal fragment (NTF) and a C-terminal fragment (CTF) occurs in many aGPCR and might offer a possible mechanism for realizing the N terminus-only/7TM-independent/*trans* function. However, our previous work showed that this cleavage is not absolutely vital for the 7TM-independent receptor mode of LAT-1, a Latrophilin homolog in the nematode *Caenorhabditis elegans* ^4^, thus indicating that other mechanisms exist. Addressing these questions regarding the details of the N terminus-only/7TM-independent/*trans* aGPCR functions is particularly difficult as their physiological implications and the receiving cells remain largely unknown. Most insights have consequently been gained from *in vitro* and *ex vivo* studies, and information in a multicellular setting on the connection and integration of the classical G protein-mediated and the 7TM-independent function is sparse and especially information is lacking on its relevance.

Here, we provide evidence that the N terminus-only/7TM-independent/*trans* function of the aGPCR Latrophilin-1 (LAT-1) in *C. elegans* is vitally involved in a wide spectrum of physiological processes within the nematode. Previous studies have shown that the receptor controls anterior-posterior cell division plane orientation in the early *C. elegans* embryo^13^ via *cis* signaling^4^ through a Gs protein cascade^14^. Further, a N terminus-only/7TM-independent function has been described in neurons^5^ and in reproduction^4^. In the current study, we show that the aGPCR regulates sperm guidance, germline apoptosis and proliferative activity of germ cells in the gonadal stem cell niche. All of these functions together cumulate in the control of reproduction and brood size. In these contexts, the absence of *lat-1* expression in the affected germ cells and gametes suggests a non-cell autonomous *trans* function exerted from adjacent somatic cells. Moreover, the absence of a G protein-mediated function indicates no simultaneous bidirectional signaling of LAT-1. Thus, we discuss the possibility of realizing isolated *trans* functions through alternative splicing of the receptor specifically generating N terminus-only variants.

## MATERIALS AND METHODS

### Materials and Reagents

All standard chemicals were from Sigma Aldrich, ThermoFisher Scientific or Carl Roth unless stated otherwise. All enzymes were obtained from New England Biolabs.

### C. elegans *maintenance and strains*

*C. elegans* strains were maintained according to standard protocols ^15^ on *E. coli* OP50 at 22 °C unless stated otherwise. Strains carrying *nIs13 [pie-1p::vab-1::GFP + unc-119(+)]; ltIs44 [pie-1p::mCherry::PH(PLC1delta1) + unc-119(+)]* were shifted to 25°C.

Wild-type worms were *C. elegans* var. Bristol strain N2 ^15^. The following alleles were obtained from the *Caenorhabditis* Genetics Center (CGC), which is funded by the NIH Office of Research Infrastructure Programs (P40 OD010440): *lat-1(ok1465)* (generated by the *C. elegans* gene knockout consortium),, *qIs153[lag-2p::MYR::GFP + ttx-3p::DsRed]* ^*16*^, *nIs13 [pie-1p::vab-1::GFP + unc-119(+)]; ltIs44 [pie-1p::mCherry::PH(PLC1delta1) + unc-119(+)]* 17, and *tnIs6 [lim-7p::GFP + rol-6(su1006)]* ^*18*^. The alleles *qaIs7524[lat-1p::GFP::lat-1 rol-6(su1006)]*, and *aprEx77[pSP5 rol-6(su1006) pBSK]* were previously generated ^4,5,13^ and *ItIs37 [pAA64; pie-1::mCherry::his-58; unc-119 (+)]* was generously provided by Diana S. Chu (San Francisco State University, USA). The following strains containing extrachromosomal or integrated arrays were generated in this study: *lat-1(ok1465) aprEx192[pTL20; pRF4; pBSK], lat-1(ok1465) aprEx193[pJL13 mCherry pBSK], lat-1(ok1465) aprEx194[pJL14 mCherry pBSK], lat-1(ok1465) aprEx216[pDM3 mCherry pBSK]*. The strain *lat-1(knu846 [lat-1 KO/KI mCherry intronic loxP::hygR::loxP])* was generated by NemaMetrix Inc. (Eugene, Oregon) using CRISPR/Cas9 genome editing. The following combination of transgenes/alleles were obtained using standard genetic techniques ^15^: *ItIs37[pAA64; pie-1::mCherry::his-58; unc-119(+)]; qaIs7524[lat-1p:GFP::lat-1 rol-6(su1006)], lat-1(knu846 [lat-1 KO/KI mCherry intronic loxP::hygR::loxP]) II; tnIs6 [lim-7p::GFP + rol-6(su1006)], lat-1(knu846 [lat-1 KO/KI mCherry intronic loxP::hygR::loxP]) II; qIs153[lag-2p::myr::GFP + ttx-3p::DsRed], lat-1(ok1465) qIs153[lag-2p::myr::GFP + ttx-3p::DsRed]*.

### *Generation of transgenic* C. elegans *lines*

All transgenic strains with stably transmitting extrachromosomal arrays were generated using DNA microinjection performed by NemaMetrix Inc. (Eugene, Oregon). Plasmids were injected at a concentration of 1 ng/μl together with the coinjection marker, a modified pPD118.33 containing *myo-2p::mCherry* (kind gift of Ralf Schnabel (Technical University Braunschweig, Germany) (30 ng/μl), and pBluescript II SK+ vector DNA (Stratagene) as stuffer DNA to achieve a final concentration of 120 ng/μl. DNA was injected into the syncytical gonad of hermaphrodites. Transgenic progeny were isolated and stable lines were selected. Multiple independent transgenic lines were established for each transgene tested.

### Brood size and lethality assay

Hermaphrodite L4 larvae were individually placed on NGM plates containing *E. coli* OP50 to lay eggs at 22 °C and transferred onto a fresh plate every 24 hours. Each day, progeny was counted until egg laying ceased. For the lethality assay, offspring were incubated at 22 °C and the number of adult/L4 animals was scored 48 hours after the mother was removed. Sterile and semi-sterile (less than 25 eggs) mothers were excluded from the assay.

### Larval development assay

To assess time of development, approximately 800 synchronized wild-type and *lat-1* L1 larvae were placed on *E. coli* OP50-seeded NGM plates and incubated at 22°C. After 8, 16, 24, 36, and 48 hours, the number of larvae in different developmental stages (L1, L2, L3, L4, adult) was scored.

### Ovulation rate assay

The ovulation rate was determined in hermaphrodites 24 hours and 96 hours post L4, respectively. Hermaphrodites were separately placed on *E. coli* OP50-seeded NGM plates and eggs/oocytes inside the uterus were counted. After 4 hours at 22 °C, eggs/oocytes on the plate and inside the uterus of the mother were counted. Ovulation rate per gonad was calculated as ([eggs in and out of uterus after 4 hours] – [initial eggs inside uterus]) / (2 * 4 hours).

### MitoTracker Red staining

To asses sperm movement *in vivo*, sperm was labelled using MitoTracker Red ^19^. L4 males were kept isolated overnight to ensure sperm accumulation. The following day, young adult males were stained with 10 µM MitoTracker Red CM-H2XRos (ThermoFisher) in M9 for 2 hours in the dark at room temperature and then left overnight to recover on plates seeded with *E. coli* OP50. The next day, 25 stained males and 25 young adult hermaphrodites, which were first anesthetized using 300 µM levamisole (Applichem), were mated on a 5 µl spot of overnight grown *E. coli* OP50. Copulation was monitored. Successfully inseminated hermaphrodites (indicated by transfer of fluorescent sperm into the uterus) were transferred separately into the wells of a 72-well plate containing 1.5 mM levamisole. Thereafter, hermaphrodites were mounted on 2% agarose pads and microscopy was conducted 1 hour post insemination. The resulting stack images of MitoTracker Red-stained sperm in the hermaphrodite uterus were evaluated as average intensity Z-projections. Correct localization of sperm was defined as proximal of the first egg next to the spermatheca, including all sperm around this egg. The percentage of correctly localized sperm was calculated as (mean fluorescence intensity of correctly localized sperm / mean fluorescence intensity of all sperm inside uterus).

### SYTO staining

Adult hermaphrodites (24 hours post L4) were washed off NGM plates with NGM lacking phosphate buffer. Worms were then incubated overnight in the dark at room temperature with 33 µM SYTO 12 Green fluorescent nucleic acid stain (ThermoFisher) in NGM and a small amount of *E. coli* OP50. To remove the stained bacteria, worms were washed three times with NGM and transferred to NGM plates with fresh *E. coli* OP50 for 1 hour. Subsequently, stained worms were anesthetized with 300 µM levamisole (Applichem) in M9 and were mounted on 2% agarose pads for microscopy.

### Antibody and DAPI staining

Antibody and DAPI staining were performed on extruded germlines. For this purpose, germlines of precisely synchronized hermaphrodites were dissected, fixed and stained as previously described ^20,21^. Adult hermaphrodites (24 hours post L4) were transferred into 300 µM levamisole (Applichem) in 0.1% Tween in PBS (PBST) for immobilization and gonad arms were exposed by cutting off the heads/tails with a scalpel blade. Gonads were fixed in 4% formaldehyde/PBST solution for 15 minutes. After washing with PBST, the specimens were incubated in 100% ice-cold methanol at -20 °C for 5 minutes.

Thereafter, the gonads were incubated overnight at 4 °C with rabbit anti-phospho Histone H3 (Ser10) (Millipore) 1:200 in 0.1% bovine serum albumine (BSA)-containing PBST. After washing three times with PBST, the gonads were incubated overnight in the dark at 4 °C with secondary goat anti-rabbit IRDye 680RD-conjugated antibody (LiCor) 1:1000 in PBST/0.1% BSA and 1 ng/µl 4,6-diamidine-2-phenylindole (DAPI, Sigma). The gonads were mounted on 2% agarose pads in Fluoromount-G (ThermoFisher) for microscopy.

To stain gonads exclusively with DAPI, fomaldehyde/methanol-fixed gonads (as for anti-PH3 staining) were incubated for 2 hours at 4 °C in the dark with 1 ng/µl DAPI. Gonads were washed three times with M9 and mounted as described above.

### *Visualizing the* lat-1p::mCherry *reporter*

Since the single copy integrated *lat-1p::mCherry* reporter used in this study shows only very faint expression, which is further reduced by even short formaldehyde fixation, we employed native microscopy to image the respective worm strains. Thereby, gonads were extruded as described above directly on a coverslip in 15 µl of 0,1% PBST containing 300 µM levamisole, which was subsequently inverted onto a glass slide and imaged immediatel

### Microscopy

All specimen were imaged using confocal imaging techniques. Differential interference contrast (DIC) and fluorescence imaging were performed using LASX software on a Leica SP8 microscope and Olympus Fluoview FV1000 software on an Olympus microscope, respectively.

Z-stacks were taken with a spacing of 0.3-2 μm, depending on the specimen (2 μm for whole worms, 0.5-1 μm for germlines, and 0.5 μm for sperm). Microscopic images were evaluated using Fiji ^22^ and ImageJ ^23^.

### Oocyte size measurements

To asses oocyte size, strains *tnIs13[pie-1p::vab-1::GFP + unc-119(+)], ltIs44[pie 1p::mCherry::PH(PLC1delta1) + unc-119(+)]*, and *lat-1; tnIs13[pie-1p::vab-1::GFP + unc-119(+)]. ltIs44[pie 1p::mCherry::PH(PLC1delta1) + unc-119(+)]* containing a cell membrane as well as a cytoplasmic marker were utilized. Worms were grown at 25°C to induce marker expression, anesthetized at young adult stage and mounted on 2% agarose pads for microscopy. Stack images of oocytes were acquired with stack borders set on the last visible part of oocyte membrane, visualized by mCherry fluorescence. Every two adjacent images were used to reconstruct part of the oocyte modelled as a truncated pyramid with the volume V = 1/3 * (slice spacing) * (A1 + A2 + sqrt(A1*A2)), with A1 and A2 being the area enclosed by the oocyte membrane in both respective images. Full oocyte volume was calculated as the sum of the volume of all truncated pyramids.

### Plexus and cap measurements of the DTC

To analyze DTC morphology, adult nematodes (24 hours post L4) expressing *qIs153[lag-2p::myr::gfp + ttx-3p::DsRed] were* anesthetized with 300 µM levamisole (Applichem) in M9 and mounted on 2% agarose pads for microscopy. A Z-projection of the resulting stacks of images was used to asses cap length (spanning the solid body of the cell) and plexus length (spanning cap and processes of the cell up to the last visible intercalating process). Measurements were based on DTC morphology described in ^16^.

### Generation of transgenes

To obtain constructs, recombineering was conducted. Accordingly, existing protocols ^24,25^ were modified as previously described ^4,13^ to generate LAT-1 transgenes using cosmids, PCR-amplified targeting cassettes and positive antibiotic selection. All transgenes are based on a construct comprising the genomic locus of *lat-1* containing *lat-1p::lat-1(1-581)::GFP* (pTL20) ^4^. To replace the promoter with tissue-specific promoters, a recombineering targeting cassette consisting of two parts, a spectinomycin selection cassette and the promoter sequence, was generated. For primer sequences, see Table S1.

#### *lag-2p::lat-1(1-581)::GFP* (pJL13)

A 2 kb promoter region of *lag-2* was amplified with primers lat1_1088F/lat1_1089R from genomic DNA of a mixed population of N2 and ligated into vector pCR2.1 using the TOPO TA Cloning Kit (ThermoFisher). From this vector the promoter was again amplified with primers lat1_1090F/ lat1_1091R with the forward primer introducing a *Pst*I site and the reverse primer containing an overhang with a homology to the vector pTL20. In parallel, the spectinomycin resistance gene was amplified from the Gateway cloning vector backbone pDON223 (Invitrogen) with primers lat1_1080F (introducing an overhang with a homology to pTL20) and lat1_1081R (containing a *Pst*I site). Both fragments were digested by *Pst*I and ligated together using a T4 DNA ligase. The ligated cassette was subsequently recombineered into pTL20 (*lat-1p::lat-1(1-581)::GFP*) replacing the *lat-1* promoter using electrocompetent SW105 cells with a heat-induced recombinase of the λ-Red recombinase system ^26^.

#### *plc-1p::lat-1(1-581)::GFP* (pJL14)

The promoter region of *plc-1* (2 kb) was obtained by amplification from genomic DNA of a mixed N2 population using primers lat1_1084F/lat1_1085R. Following ligation into vector pCR2.1 using the TOPO TA Cloning Kit (ThermoFisher), the promoter was amplified with primers lat1_1086F/ lat1_1087R. While the forward primer introduced a *Pst*I site, the reverse primer contained an overhang with a homology to the vector pTL20. In parallel, the sequence of the spectinomycin resistance gene was amplified from the Gateway cloning vector backbone pDON223 (Invitrogen) with primers lat-1_1080F (containing an overhang with a homology to pTL20) and lat-1_1081R (introducing a *Pst*I site). Both fragments were digested by *Pst*I, ligated with a T4 DNA ligase and the resulting fragment was recombineered into pTL20 (*lat-1p::lat-1(1-581)::GFP*) replacing the *lat-1* promoter using electrocompetent SW105 cells with heat-induced recombinase of the λ-Red recombinase system ^26^.

#### *lim-7p::lat-1(1-581)::GFP* (pDM3)

The promoter region of *lim-7* (2 kb) was retrieved by PCR with primers lat1_1082F (containing a *Pst*I site) and lat1_1083R (harbouring a homology sequence to pTL20) from vector pOH323, which was a kind gift from Oliver Hobert (Columbia University, New York, USA) ^18^. In parallel, primers lat1_1080F (containing an overhang with homologies to pTL20) and lat1_1081R (introducing a *Pst*I site) were used to amplify the spectinomycin resistance gene from the Gateway cloning vector pDON223 (Invitrogen). Both PCR products were digested with *Pst*I (NEB) and ligated using a T4 DNA ligase. The resulting fragment was recombineered into pTL20 (*lat-1p::lat-1(1-581)::GFP*) replacing the *lat-1* promoter using electrocompetent SW105 cells with heat-induced recombinase of the λ-Red recombinase system ^26^.

### RNA extraction and rapid amplification of 3’ cDNA ends with PCR (RACE-PCR)

Total RNA was extracted from a mixed population of wild-type hermaphrodites using TRI Reagent (Sigma) according to the manufacturer’s protocol and were transcribed into cDNA with the 2^nd^ generation 5’/3’ RACE Kit (Roche) in combination with the Expand High Fidelity PCR System (Roche) as described in the supplier’s instructions. To ensure cDNA synthesis from mainly mRNA, a specific oligo(dT)-anchor primer was used. The cDNA mix was directly utilized as a template for the RACE-PCR. Amplification was run with a PCR anchor primer corresponding to the sequence of parts of the oligo(dT)-anchor primer used for cDNA synthesis and the gene-specific primers lat1_1365F and lat1_1394F, respectively. PCR products were separated via gel electrophoresis, cut out, purified with the Wizard SV Gel and PCR Clean-Up System (Promega) and cloned into the vector pCR2.1 using the TOPO TA Cloning Kit (ThermoFisher) for further analysis. For primer sequences, see Table S1.

### Transcript variant analyses

Publicly available RNA-Seq datasets from adult wild-type hermaphrodite *C. elegans* var. Bristol N2 were downloaded via the Sequence Read Archive ^27,28^, reads were aligned to the WBcel235 genome ^29^ using STAR ^30^ and transcript assembly was performed using StringTie ^31^. Full-length transcripts were inspected and further analyses including visualization were conducted with the Integrated Genomics Viewer ^32^. Accession numbers and mapping statistics are given in Table S2.

### Statistical analysis

Numerical assay data were analyzed with GraphPad Prism version 9.2.0 (GraphPad Software). If not stated otherwise, the data are presented in box plots with 90% confidence interval. Statistical analyses were performed using a two-tailed Student’s t-test for comparing two groups and a two-way ANOVA in combination with a Bonferroni post-hoc test for comparing multiple groups, respectively. P values of p ≤ 0.05 were considered statistically significant. All details are given in the respective figure legends.

## RESULTS

### LAT-1 controls sperm movement in hermaphrodites

Previously, we showed that the membrane-tethered LAT-1 N terminus (aa 1-581, termed LNT) is sufficient to ameliorate the fertility defect in *C. elegans* homozygous for the null allele *lat-1(ok1465)* (hereafter referred to as *lat-1*) ^4,13^, indicating that the unusual 7TM-independent function of the receptor is involved in controlling aspects of reproduction. To address the question whether this is caused by a single LAT-1 function or several ones in different processes cumulating into the regulation of reproduction, we delineated the receptor’s physiological impact by dissecting the *lat-1* mutant defects greater detail. These are generally characterized by a reduced brood size (wild-type: 228.2 ± 3.6; *lat-1*: 117.4 ± 4.0) ^4,13^. At least three distinct processes involving the 7TM-independent LAT-1 function were identified in the self-fertilizing *C. elegans* hermaphrodite gonad, which produces a fixed number of sperm during larval development that are subsequently stored in the spermatheca, and variable amounts of oocytes in adulthood (Fig. 1).

**Figure 1.**
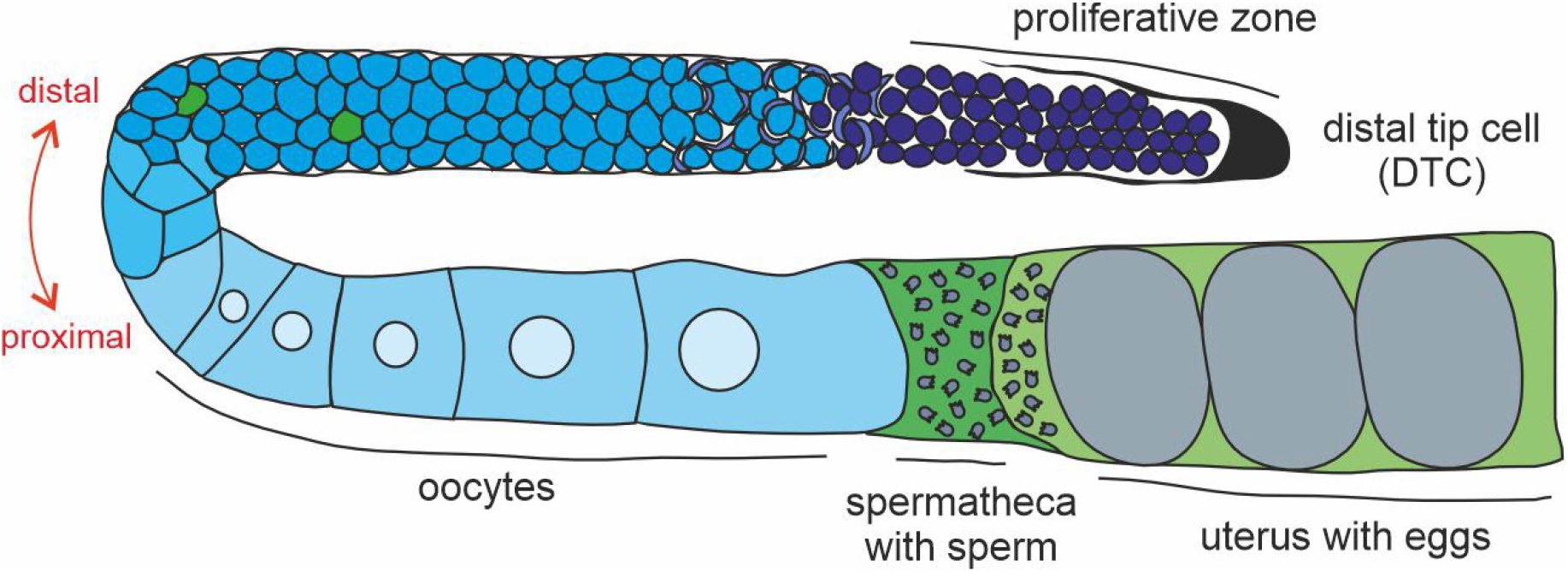
The *C. elegans* gonad. Shown is one of the two symmetrical U-shaped gonad arms of a *C. elegans* adult hermaphrodite. In the fourth larval stage, a fixed number of sperm (approximately 150 per gonad) is produced and stored inside the spermatheca. Subsequently, the gonad switches to continuous oocyte production only. In the distal gonad, germ cell nuclei are not surrounded by a complete membrane but reside in a common cytoplasm. They continually self-renew by mitotic division in the proliferative zone (dark blue), which is distally enclosed by the distal tip cell (DTC) plexus. Subsequently, germ cells enter meiotic divisions (crescent-shaped nuclei) and progress further through meiosis, where some germ cells undergo apoptosis (marked in green). Here, the cells are surrounded by the gonadal sheath cells (not shown). Near the loop, nuclei start to cellularize and to obtain plasma membranes. Due to coordinated contraction of the gonadal sheath cells, oocytes are pushed into the spermatheca, fertilized, and eggs are flushed into the uterus. After completing the first set of embryonic divisions, they are laid through the vulva (not shown).

We first analyzed *lat-1* mutant sperm as previously obtained data led to the hypothesis that LAT-1 plays some role in sperm function ^4^. Mutant hermaphrodites produced approximately the same amount of sperm as wild-type individuals (Fig. 2A, B) and the number of anucleate residual bodies, which are formed during spermatogenesis as remainders of budding spermatids (summarized in ^33^), appeared to be constant (Fig. 2C, D). These data indicate largely intact sperm development and the correct timing of the sperm-oocyte switch during the transition of the last larval stage into adulthood. As *lat-1* nematodes lay overall fewer fertilized eggs ^4^ and thus, use less sperm than wild-type worms in a given time period, this should result in an accumulation of residual sperm over time. To assess a possible decline in sperm amount during the fertilization period resulting the reduced number of offspring, we examined sperm in the spermatheca 48 hours after the first ovulation (Fig. 2E) together with the course of egg-laying (Fig. 2F). Sperm amounts in the spermatheca 48 hours after the first ovulation was slightly, albeit not significantly reduced compared to the amount in wild-types. At the same time, significantly less sperm was used for fertilization indicating a loss of sperm during adulthood. Sperm loss was evaluated by subtracting the number of laid eggs and residual sperm within the spermatheca from the initial sperm count (Fig. 2E). As sperm loss can be caused by impaired motility or defective directed locomotion of sperm, which leads to them being flushed out during egg-laying (summarized in ^34^), our data suggest that LAT-1 could play a role in sperm movement. Consistent with this hypothesis, the ovulation rate in *lat-1* mutants, which was initially similar to the one of wild-type worms, declined more rapidly over time (Fig. 2G). This is plausible as in *C. elegans*, ovulation is stimulated by signals produced in spermatozoa (reviewed in ^34^). Both phenotypes were rescued by the transgenic LNT (Fig. 2E, G), showing that the role in sperm movement is mediated by the 7TM-independent function of LAT-1. It has to be noted that worms were precisely staged in all experiments, also to account for the fact that some *lat-1* mutants are developmentally slower than wild-type controls ^13^ (Fig. S1).

**Figure 2.**
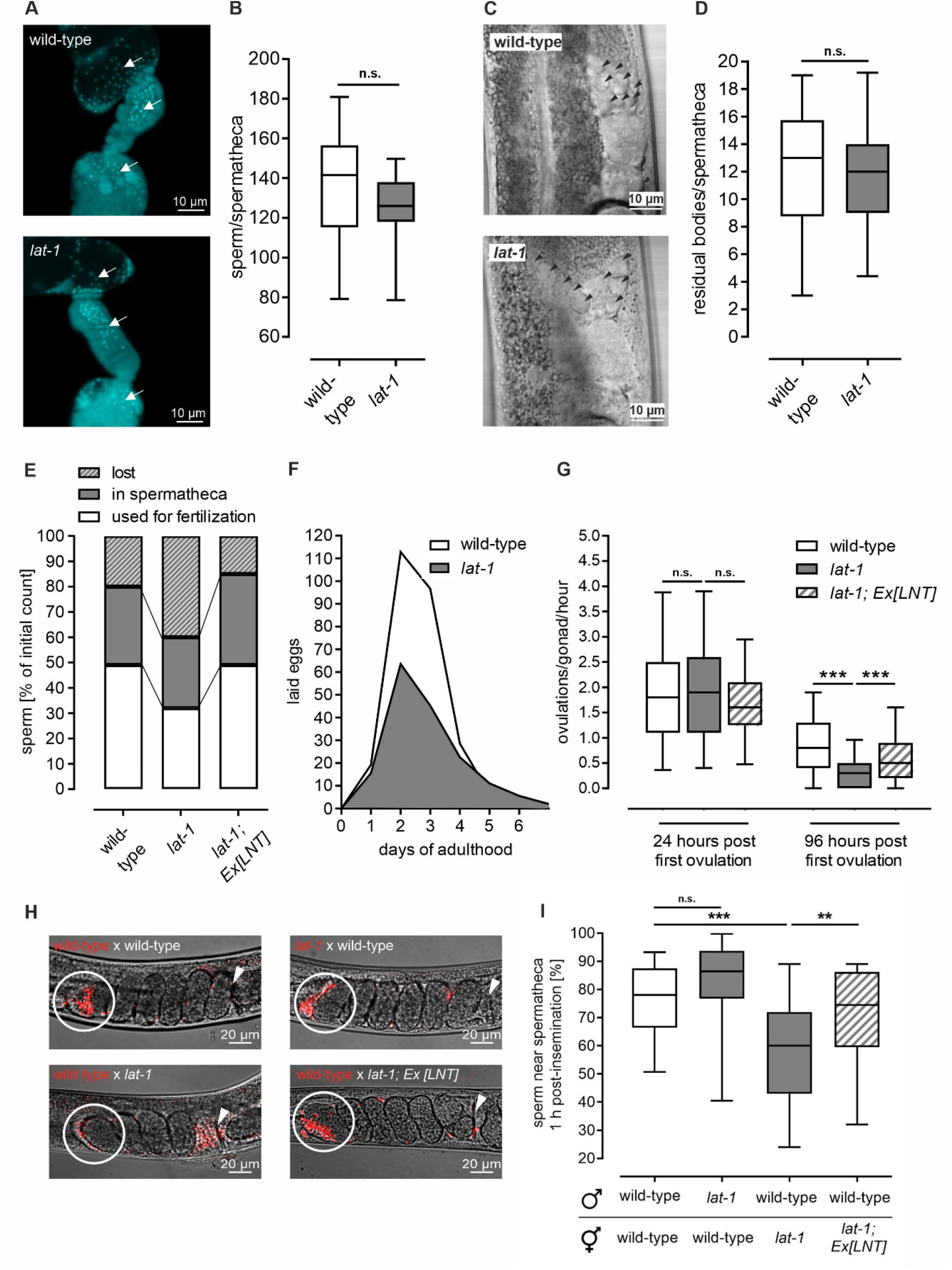
Nematodes lacking *lat-1* display impaired sperm movement. (A) Spermathecae of L4 + 1 day *lat-1* mutant hermaphrodites contain the same amount of sperm (white arrows) as wild-type specimen. Gonads of adult hermaphrodites were dissected and subsequently stained with DAPI. (B) Quantification of DAPI stainings as shown in (A) reveals that the number of sperm inside the spermatheca is indifferent from the one in wild-type nematodes. Data are shown as box plots with 90% confidence interval of 4 independent experiments, n ≥ 20. n.s. = not significant. (C) Residual bodies (black arrow heads) located close to the spermatheca in *lat-1* mutant and wild-type gonads. Shown are DIC images of unstained living L4 + 1 day-old hermaphrodites. (D) Quantification of anucleate residual bodies based on DIC images as displayed in (C) revealed no significant difference in number between wild-type nematodes and *lat-1* mutants. Data are shown as box plots with 90% confidence interval of 4 independent experiments, n ≥ 24. n.s. = not significant. (E) Sperm amount 48 hours post the first ovulation. Sperm was DAPI-stained and sperm loss was calculated as follows: ((initial sperm in spermatheca) - (sperm in spermatheca after 48 hours) - (eggs laid in 48 hours)) / (initial sperm in spermatheca). This loss is ameliorated in transgenic lines expressing LNT. It has to be noted that due to experimental limitations, the initial sperm count in the spermatheca had to be taken from different animals than the count after 48 hours. n ≥ 15. (F) Hermaphrodites lacking *lat-1* exhibit a continuously smaller brood size over their entire reproductive period with the maximal reduction being visible on the second and third day of egg laying. n ≥ 75 in 9 independent replicates. (G) The ovulation rate in *lat-1* mutants decreases faster than in wild-type individuals or in *lat-1* mutants expressing the LNT. The ovulation rate per gonad per hour was calculated as follows: ((eggs in and out of uterus after 4 hours) -(initial eggs inside uterus)) / (2 * 4 hours). Data are shown as box plots with 90% confidence interval of 4 independent experiments, n ≥ 51. n.s. = not significant; *** p < 0.001. (H) Distribution of male *lat-1* and wild-type sperm, respectively, after mating to hermaphrodites of different genotypes. The motility of *lat-1* sperm after mating with wild-type hermaphrodites appears to be intact, as indicated by correct localization near the spermatheca (white circle) 1 hour post insemination. Conversely, wild-type sperm inside *lat-1* hermaphrodites do not reliably localize next to the spermatheca, which can be rescued by transgenic complementation of the LAT-1 N terminus (LNT). Mating assays were performed with wild-type and *lat-1* mutant nematodes in different constellations. Sperm was stained with MitoTracker Red by incubation of living young adult males prior to the mating and sperm location was monitored 1 hour after mating occurred. (I) Sperm of wild-type males do not properly migrate to the spermatheca in *lat-1* mutant hermaphrodites after mating while sperm of *lat-1* mutant males show no impaired movement towards the spermatheca of wild-type hermaphrodites. Quantification was performed from images as shown in (H). Data was acquired in at least 3 independent experiments with n > 20 biological replicates. n.s. = not significant; ** p < 0.01; *** p < 0.001.

To further assess the hypothesis that LAT-1 is essential for sperm movement, we monitored MitoTracker Red-labelled sperm after mating of males and hermaphrodites of different genotypes. Sperm of *lat-1* mutant males did not show impaired movement towards the spermatheca of wild-type hermaphrodites, indicating largely intact sperm motility (Fig. 2H, I). In contrast, wild-type sperm inside *lat-1* hermaphrodites did not localize near the spermatheca to the same extent as in the respective wild-type control. This defect was ameliorated by transgenic expression of the LNT in the hermaphrodite. Thus, our data indicate a defect in sperm guidance originating from *lat-1* hermaphrodites rather than impaired sperm motility.

### LAT-1 modulates germ cell apoptosis and oocyte quality

Besides the effect on sperm movement, we found that secondly, the 7TM-independent mode of LAT-1 also affects oocytes. We observed an increase of germ cell apoptosis by SYTO 12-staining of *lat-1* mutant hermaphrodites, which was rescued by transgenic expression of the *LNT* (Fig. 3A, B). Germ cells normally form oocytes and it is physiological process during *C. elegans* oogenesis that some of them undergo apoptosis (Fig. 1) (summarized in ^35^). While its role is debated, one possible function is to provide nutrients for developing oocytes ^36,37^. Oocytes gaining too little nutrients are of inferior quality, reflected by reduced oocyte size ^37^ and increased embryonic lethality ^38^. Surprisingly, *lat-1* oocytes displayed both of these characteristics (Fig. 3C-E). As nematodes null for *lat-1* generally show embryonic lethality due to the receptor’s function in early development ^13,14^, it is impossible to assess oocyte quality by directly quantifying embryonic lethality. Thus, we investigated *lat-1* worms expressing the *LNT*. As this construct only rescues fertility defects ^4^, any lethality observed in the corresponding strain should be attributed to LAT-1 function during embryogenesis. Comparison with *lat-1* mutants revealed that embryonic lethality is less pronounced in the presence of the LNT (Fig. 3E), indicating a cause other than developmental defects such as inferior oocyte quality. These data point towards *lat-1* oocytes gaining too little nutrients, which can imply insufficient apoptotic events rather than too many. Therefore, *lat-1* worms may not display a genuine increase in apoptosis but have other defects rendering apoptotic cell accumulation.

**Figure 3.**
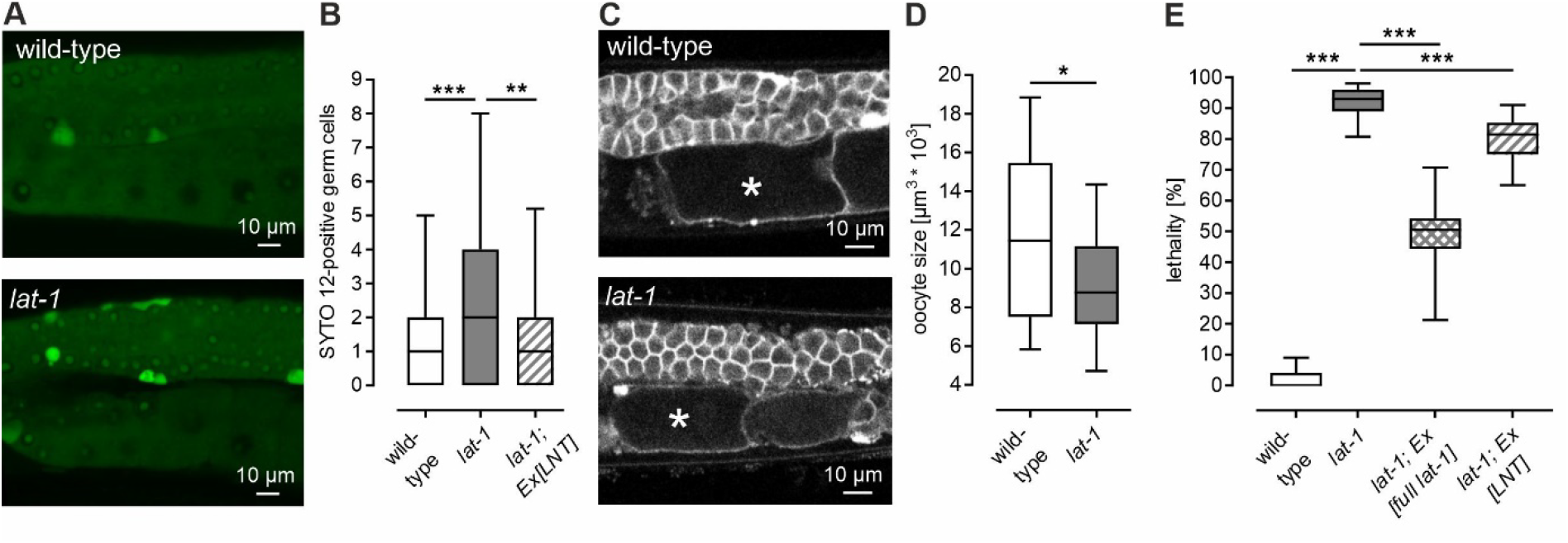
In the absence of LAT-1 function, germ cell apoptosis is increased. (A) Germlines of *lat-1* mutants exhibit more SYTO 12-stained germ cells compared to wild-type controls. Living adult nematodes 24 hours post L4 were stained with SYTO 12 as a marker for apoptotic cells. (B) Quantification of SYTO 12 stainings from (A). Data are shown as box plots with 90% confidence interval, n ≥ 33. ** p < 0.01; *** p < 0.001. (C) Oocytes (marked with an asterisk) are significantly smaller in *lat-1* mutants than in wild-type individuals. Oocyte membranes were visualized in living young adult hermaphrodites by expression of the construct *pie-1p::mCherry::PH(PLC1delta1)*. (D) Quantification of (C) confirms that oocyte size is decreased in *lat-1* mutants. Data are shown as box plots with 90% confidence interval, n ≥ 15. * p < 0.05. (E) The LNT rescues embryonic lethality of *lat-1* mutants to a small but significant extent while amelioration by the full-length *lat-1* construct is much stronger and is considered the maximum rescue that can be reached. Assays were performed in 4 independent experiments (n ≥ 30) and data are shown as box plot with 90% confidence interval. *** p < 0.001.

### LAT-1 is involved in germ cell proliferation

A third function of LAT-1 was identified in the proliferation of germ cells leading to mature oocytes. The generation of germ cells during the adult life of *C. elegans* hermaphrodites is a tightly regulated process, which entails the balance between cell proliferation in its gonadal stem cell niche and differentiation into germ cell progenitors (Fig. 1). In *lat-1* mutant hermaphrodites, germ cell proliferation was reduced. We observed a decrease in mitotic events indicated by a smaller number of nuclei stained with an anti-phospho-histone H3 (Ser10) (PH3) antibody, which marks the mitotic (M) phase (Fig. 4A). Transgenic complementation with the *LNT* ameliorated this defect at least partially (Fig. 4B), suggesting that this LAT-1 function is also based on the 7TM-independent mode.

**Figure 4.**
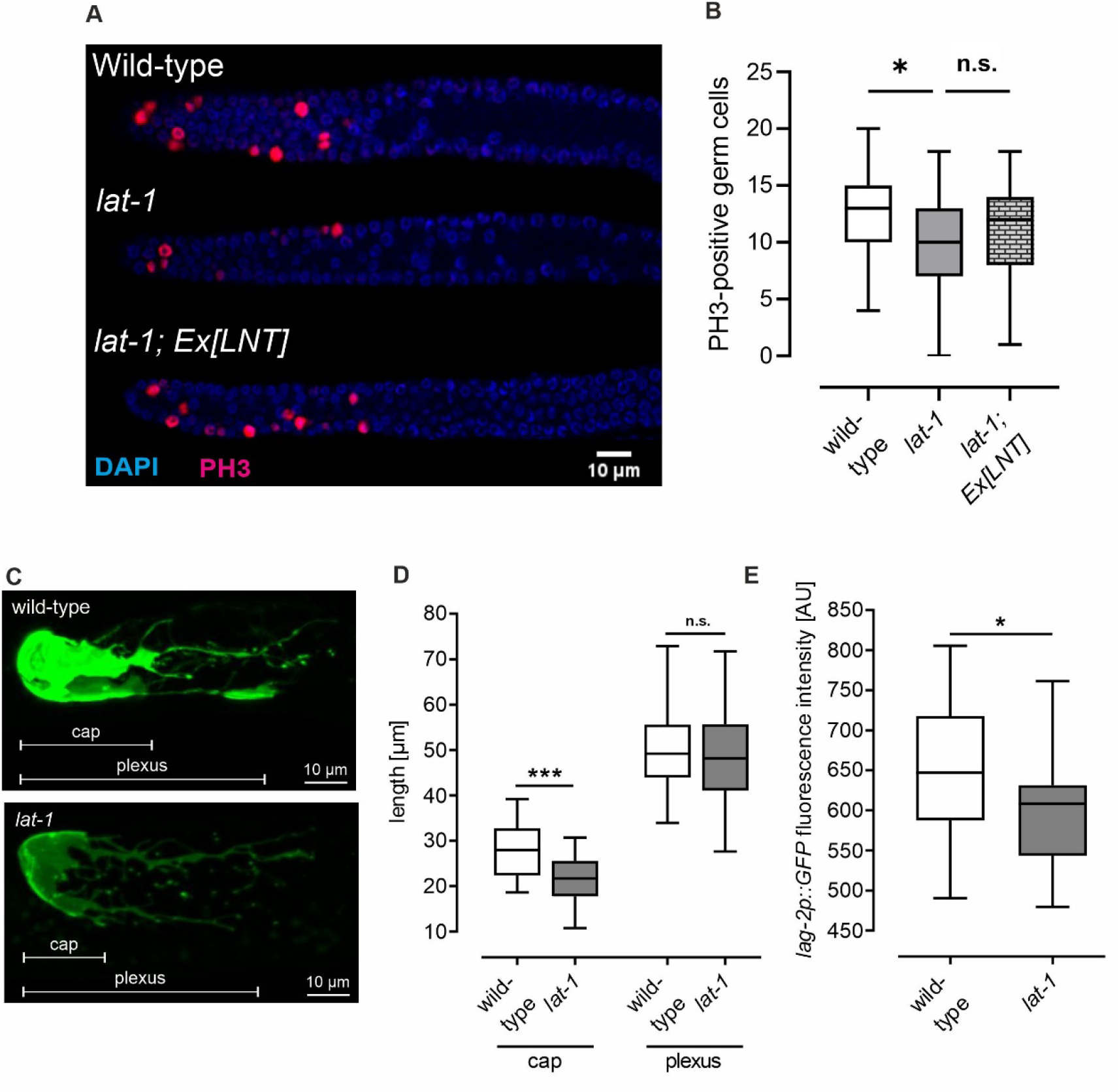
*lat-1*-deficient animals show defects in germ cell proliferation. (A) M phase nuclei are decreased in *lat-1* hermaphrodites, which can be ameliorated by *LNT* expression. Gonads were dissected and DAPI-(blue) and PH3-(magenta) stained to visualize all and M phase nuclei, respectively. (B) Quantification of M phase nuclei. n ≥ 19 n.s. = not significant; * p < 0.05. (C) Wild-type and *lat-1* mutant DTC visualized by *lag-2p::myr::GFP* expression in 24 hours post L4 animals. The morphology of the *lat-1* mutant DTC is largely intact compared to the wild-type DTC. Loss of *lat-1* only leads to a slightly reduced cap length, while the extension of the plexus over the proliferative zone is unchanged. Marker expression appears fainter in *lat-1* mutants. (D) Quantification of DTC parameters shown in (C) confirm the reduced cap length of the DTC in *lat-1* mutants compared to wild-type controls. Data are shown as box plots with 90% confidence interval, n ≥ 42. n.s. = not significant; *** p < 0.001. (E) Quantification of expression levels shown in (D). Boxplots with 90% confidence interval, n ≥ 34. * p < 0.05.

The main source of the signal controlling cell proliferation in the proliferative region is known to be the DTC ^39^. Thus, we elucidated the morphology of this cell in precisely staged 24 hours-post L4 *lat-1* hermaphrodites to exclude that a malformed DTC was the cause for the observed defect. This is especially important as malformations and developmental defects are known to generally occur in *lat-1* mutants ^13^. While the plexus length was indistinguishable from that of wild-type nematodes, the cap length was slightly reduced (Fig. 4C, D) indicating that DTC morphology is largely normal and that the observed proliferation defects are likely not caused by DTC malformation. The analyses were performed using a *lag-2* promoter-driven GFP as a marker specific for the DTC. LAG-2 is the ligand of the Notch receptor GLP-1, which is the major player in the regulation of germ cell proliferation and differentiation. Interestingly, the absence of LAT-1 leads to a reduced expression of *lag-2* indicated by the decreased fluorescence of the *lag-2p::GFP* construct (Fig. 4E), suggesting an effect of LAT-1 on Notch pathway components.

### LAT-1 elicits its functions from the somatic gonad in *trans*

All three identified functions of LAT-1, in sperm guidance, germ cell apoptosis/oocyte quality and germ cell proliferation, have been shown to be 7TM-independent functions of the aGPCR. To gain insights into the mechanisms underlying these functions, we addressed the question, from which cells the LAT-1 function originates. Thus, we first generated a detailed expression profile of the receptor in the hermaphrodite gonad. Utilizing an integrated transcriptional *lat-1p::GFP*, we observed strong *lat-1* expression in the entire somatic gonad of adult hermaphrodites, including the distal tip cell (DTC), the gonadal sheath cells and the spermatheca, but not in germ cells (Fig. 5A-C). To verify that the lack of germline expression was not due to silencing effects of the used transgenes, a single-copy integrated transcriptional *lat-1p::mCherry* was employed, which was generated using CRISPR/Cas9 genome editing. Like the other constructs, this single-copy integrated *lat-1p::mCherry* also showed expression of *lat-1* in the gonadal sheath cells (Fig. 5D) and the DTC (Fig. 5E), but not in germ cells.

**Figure 5.**
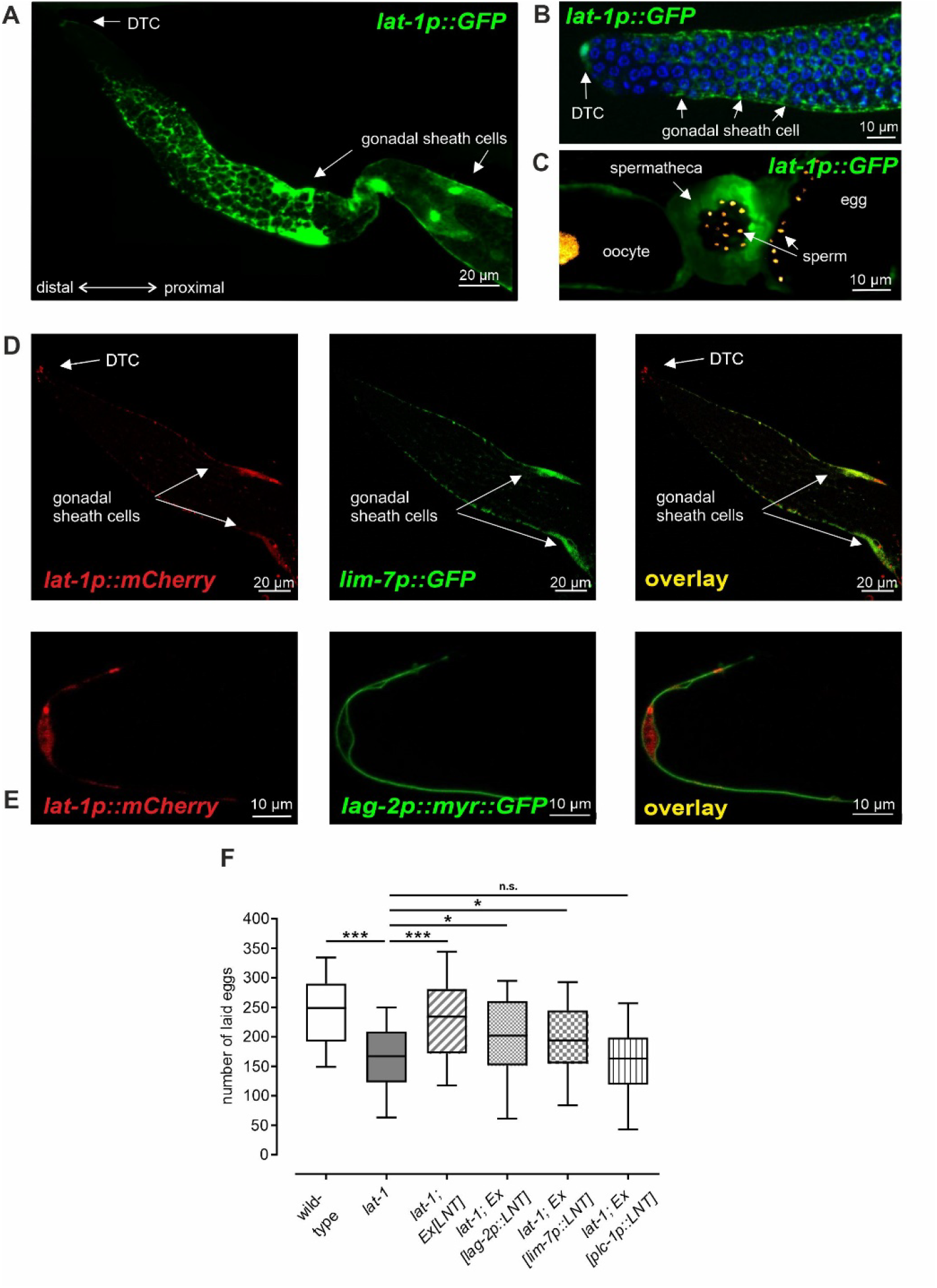
The LAT-1 N terminus acts from somatic cells of the *C. elegans* gonad in a non-cell autonomous manner. (A)-(E) Expression of *lat-1* in the hermaphrodite gonad. The receptor is present in all somatic cells of the gonad: distal tip cell (DTC), gonadal sheath cells and spermatheca. However, no expression is detectable inside germ cells, as well as in fully differentiated sperm or oocytes. For visualizing *lat-1* expression, either a multi-copy integrated *lat-1p::GFP* (A)-(C) or a CRISPR/Cas9 genome-edited single-copy integrated *lat-1p::mCherry* (D), (E) was employed. Germ cell nuclei (C) were labeled by expressing *pie-1::mCherry::his-58*, gonadal sheath cells (D) by expressing *lim-7p::GFP* and DTC (E) by using *lag-2p::myr::GFP*. (F) Tissue-specific expression of the LAT-1 N terminus (LNT) using promoters with activity restricted to distinct cell types in the somatic gonad reveals in which location LAT-1 is required to fulfil specific functions. (F) While expression of LNT in the distal tip cell (*lag-2p::LNT*) and the gonadal sheath cells (*lim-7p::LNT*) generally results in an amelioration of the brood size defect, sole expression in the spermatheca (*plc-1p::LNT*) does not. Data are given in box plots with 90% confidence interval of 5 independent experiments. n ≥ 34. n.s. = not significant; * p < 0.05; *** p < 0.001.

To test whether the somatic gonad really is the origin of the LAT-1 7TM-independent function, we specifically expressed the *LNT* in the DTC (*lag-2p::LNT*), the gonadal sheath cells (*lim-7p::LNT*), and the spermatheca (*plc-1p::LNT*) of *lat-1* mutants, respectively. Subsequently, the brood size was quantified to assess whether the LNT ameliorates the fertility defects in general. While expression in the DTC and gonadal sheath cells rescued at least partially (to a lesser extent than the LNT driven by its endogenous promoter), expressing in the spermatheca did not (Fig. 5F). These data highlight the fact that the LAT-1 7TM-independent function is required in the somatic gonad to control reproduction in a non-cell autonomous manner.

### The LAT-1 transcript variant repertoire contains N terminus-only receptor versions

As the LAT-1 7TM-independent functions act in a completely different biological context than that of its 7TM-dependent signal in embryogenesis ^4,13^, the question arose as to why such a large receptor molecule is produced only when the extracellular entity is mediating a function in distinct contexts. The release of the N terminus by cleavage of the aGPCR at the GPS, the motif in the GAIN domain capable of autoproteolysis, could be one mechanism. However, our previous findings indicate that cleavage is not essential for overall LAT-1 function ^4^. Interestingly, a recent study suggests the presence of several premature poly-A sites within *lat-1* transcripts that, among others, render LAT-1 molecules consisting only of the N terminus ^40^. To gain insights into the prevalence of such variants, we analyzed existing transcriptome data generated by RNA-Seq from day 1 adult wild-type hermaphrodites ^41^, in which the coverage was sufficient for splice variant analyses (Fig. 6A). Generally, with more than 20 transcript forms, the repertoire of *lat-1* variants can be considered extensive. Although the largest fraction consisted of the entire receptor molecule (exons 1-8), approximately 2.3% of the variants comprised only the N-terminal exons in different combinations and versions, which are potentially able to transmit the 7TM-independent function (Fig. 6B). A 3’ rapid amplification of cDNA-ends with PCR (RACE-PCR) from the mRNA of wild-type hermaphrodites confirmed the general presence of N terminus-only variants (Fig. 6C), suggesting that these variants of *lat-1* exist *in vivo*. It has to be noted that, to ensure correct transcript composition, rescue analyses were always performed with constructs containing the genomic locus of *lat-1* or modifications of it.

**Figure 6.**
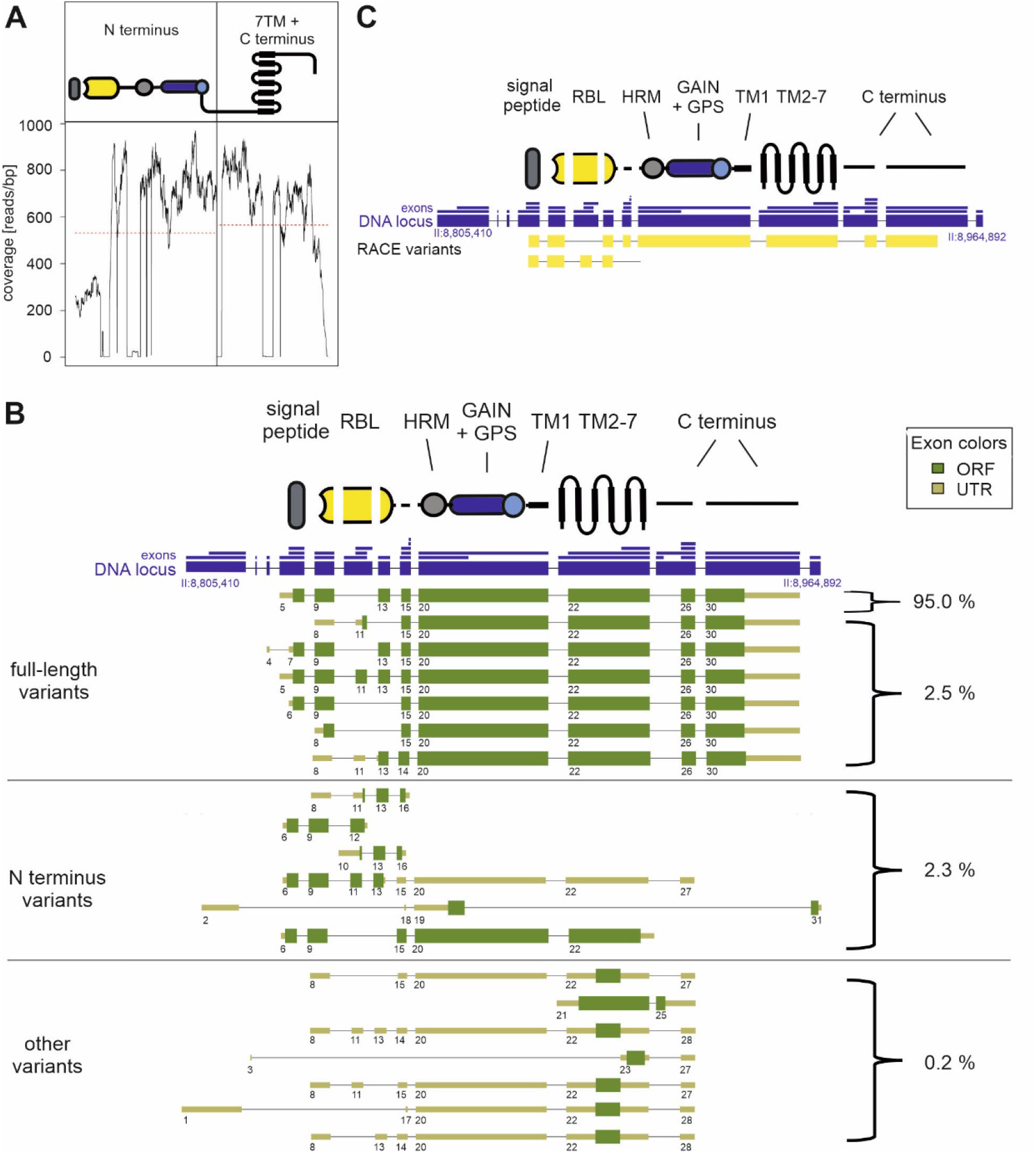
Several transcripts containing only the N terminus of LAT-1 exist. (A) Read coverage of the *lat-1* locus extracted from available RNA-Seq data of day 1 adult wild-type hermaphrodites ^41^ shows an almost uniform distribution of reads over the entire locus. (B) Transcript variant repertoire of *lat-1* identified from RNA-Seq data ^41^. Several full-length variants exist which mostly differ in their N-terminus composition. N terminus only-containing variants make up 2.3% of the total *lat-1* variants in the hermaphrodite and seem mostly not to be membrane-anchored. Other variants, which lack the N terminus, constitute less than 1%. Only full-length variants with an incidence of more than 0.01% are depicted. The genomic locus of LAT-1 is shown with its longest exons (large blue boxes) and size-condensed introns (faint blue lines). All exons found in the analysis are separately plotted above the locus (small blue boxes). The individual exon arrangements of transcripts are shown numbered. Transcripts were defined as a numeric sequence of exons. The longest bona fide open reading frame (ORF) is depicted in thick green boxes while the non-protein coding 5′ and 3′ untranslated regions (UTRs) are displayed thinner and in light green. 3′ end exons with minor differences in length but identical 5′ splice acceptor sites are considered as one 3′ end exon. Different composition of the 5′ start exon, 3′ end exon and/or exons are considered as individual variants. The exact positions of the exons forming the variants are given in Table S3. (C) 3’ RACE analyses of wild-type hermaphrodites rendered among others the full-length variant, which has been shown to be the most abundant one in RNA-Seq analyses. Further, a variant comprising the N terminus including RBL and HRM domain, but not the GAIN domain was amplified. RBL = rhamnose-binding lectin domain, HRM = hormone-binding domain, GPS = GPCR proteolytic site, GAIN = GPCR autoproteolysis-inducing domain.

## DISCUSSION

It is becoming increasingly clear that the highly unusual dual mode of classical *cis* signaling via G proteins and *trans* functions mediated solely through the N terminus (independent of 7TM and C terminus) are inherent to several aGPCR ^4,6-8,13^. While the *cis* mode is relatively well studied in many cases, knowledge on the molecular details as well as the impact of the *trans* function is sparse, limiting our understanding of how such functions are integrated in multicellular settings. In the present study, we show that the *trans*/7TM-independent mode is a frequently employed concept involved in different physiological and cellular contexts and that, in contrast to other aGPCR ^6,8^, it does not seem to engage in simultaneous bidirectional signaling of the receptor. We identified three contexts of N-terminus activity in the *C. elegans* reproductive system, jointly balancing the brood size of the nematode. Thereby, the LAT-1 7TM-independent/*trans* mode of action affects both gamete types and their function with seemingly diverse effects on sperm movement, germ cell apoptosis, and proliferation.

Although it does not appear to be involved in sperm development, the LAT-1 *trans* function seems to be essential for sperm locomotion. The corresponding defect caused by the absence of the receptor is characterized by sperm loss occurring in the course of *lat-1* mutant adulthood (Fig. 2). Impaired sperm movement can lead to sperm being flushed out of the reproductive system during egg laying (reviewed in ^34^). Possible reasons for this can generally be faulty sperm guidance or defective sperm, as previously hypothesized ^4^. Our analyses strongly suggest LAT-1 to be involved in the sperm guidance signaling machinery of the hermaphrodite, thereby affecting sperm movement.

We also found increased numbers of apoptotic cells in the proximal part of *lat-1* mutant germlines indicating a role of the receptor in programmed cell death (Fig. 3). Future analyses will need to clarify whether this is due to an increased rate of apoptosis itself or defective clearance of apoptotic germ cell corpses. A function of LAT-1 in the regulation of apoptosis is intriguing since the mouse Latrophilin LPHN1/ADGRL1 ^42^ as well as some other aGPCR ^43,44^ have previously been associated with programmed cell death.

Our data further show that the LAT-1 7TM-independent/*trans* function modulates cell proliferation (Fig. 4). The observed decrease in PH3-positive germ cells bears resemblance to mutants of the Notch pathway, which governs the balance between germ cell proliferation and entry into meiosis ^39,45-49^. Transcript analyses of the aGPCR Gpr126/ADGR in Notch-deficient mice^50^ gave a preliminary indication that aGPCR might indeed be candidates for cross-talk with Notch signaling. However, no receptor has yet been identified as a component or direct regulator of this essential and highly conserved pathway and thus, the hypothesis of an aGPCR-Notch pathway interaction remains speculative. Intriguingly, *lat-1* mutants show a decreased expression of the Notch ligand *lag-2* (Fig. 4), suggesting a transcriptional regulation. However, in the context of LAT-1 regulating proliferation, the receptor is able to function without its 7TM, rendering it unable to target transcriptional pathways via canonical signaling. It would therefore be intriguing to investigate whether the LAT-1 N terminus can affect the Notch pathway in *trans*, thereby regulating proliferation in the stem cell niche of the *C. elegans* germline.

Since the three functions of LAT-1 affect three distinct areas of the germline, but *lat-1* is expressed only in the surrounding somatic cells, it is likely that the receptor functions are exerted non-cell autonomously and thus, in *trans*. However, it remains to be determined whether the three functions employ similar mechanisms on a molecular level. Given the plethora of physiological implications of LAT-1 identified in this study and previously ^4,5^, it is conceivable that several more LAT-1 functions exist, and although we did not detect any germline expression with our constructs, LAT-1 presence in the germline cannot be ruled out entirely. Despite the clear evidence that several aGPCR are able to engage in 7TM-independent/*trans* modes of action ^4,6-9^, it is still debated whether these functions are signaling or if they involve adhesion or even other modes of action. While adhesive components of LAT-1 cannot be excluded, the almost unchanged morphology of *lat-1*-expressing cells and their surrounding tissues in the absence of the receptor (Figs. 4, 5) make a sole function as a structural component of cell-cell adhesion complexes unlikely. The potential role of the slightly reduced DTC cap size of *lat-1* mutants in this context remains elusive and requires further investigation.

Intriguingly, the *trans* function seems to be separable at a molecular level from the *cis* function due to alternative splicing, giving rise to a multitude of transcript variants (Fig. 6). Thereby, isolated N-terminal variants can be expressed *in vivo*, and it is plausible that they are regulated separately from the full-length receptor. These data are also in concordance with recent studies on other aGPCR showing a huge tissue-specific transcriptional variability as a potential common principle in the regulation of aGPCR function ^51-53^. It has to be noted that no information is available on the presence of N-terminal variants in the *C. elegans* somatic gonad cells, only data on whole adult hermaphrodites are accessible. The small proportion of N-terminal variants (2.3%) might be due to this heterogeneity of the samples. Besides different transcript variants, it is also conceivable that the 7TM-independent function is brought about by the liberation of the N terminus through autoproteolysis at the GPS. This mechanism, which is a hallmark feature of many aGPCR, has been described for LAT-1, but in previous studies we found that cleavage does not have a major impact on receptor function ^4^. It is known that LAT-1 *cis* function is essential for early embryonic development ^13^. While *cis* and *trans* function seem to be involved in different developmental stages, it has to be noted that future analyses can unravel LAT-1 *cis* effects in adult nematodes and *trans* functions in the embryo.

In summary, we show that the LAT-1 N terminus acts in *trans* in multiple, distinct physiological settings, underlining the importance as well as the versatility of this mode of action. We further demonstrate that next to the possibility of liberating the extracellular domains of aGPCR via autoproteolysis, alternative splicing can generate receptor N termini to exert 7TM-independent *trans* functions. It would be highly interesting to see whether the characteristics of *trans* signaling found for the prototypic aGPCR LAT-1 in this study are conserved among species and are moreover applicable to the entire class of aGPCR.

## ACKNOWLEDGEMENTS

The authors thank Jana Winkler for help with establishing fertility phenotyping assays, Sonja Kallendrusch for advice and support with microscopy, and Samantha Hughes for helpful discussions. We are very grateful to Oliver Hobert and Ralf Schnabel for kindly providing plasmids.

This work was supported by grants from the Deutsche Forschungsgemeinschaft (DFG, German Research Foundation) through CRC 1423 (project number 421152132; C04 (S.P., T.S.)), FOR 2149 (project number 246212759; P02 (S.P.) and P04 (T.S.)), Pr1534/1-1 (project number 254080357 (S.P.)), the European Social Fund (S.P.) and the Federal Ministry of Education and Research (BMBF), Germany (A.B.K., FKZ: 01EO1501).

## AUTHOR CONTRIBUTIONS

S.P. conceived and designed the study and the experimental approaches. D.M., V.E.G., F.F., J.L.S., and A.B.K. carried out the experiments. D.M., V.E.G., W.B.P., A.B.K., T.S., and S.P. analyzed the data. D.M., V.E.G, W.B.P, and S.P. wrote the manuscript.

