## Supplemental Information for "The N terminus-only (*trans*) function of the Adhesion GPCR Latrophilin-1 controls multiple processes in reproduction of *C. elegans*"

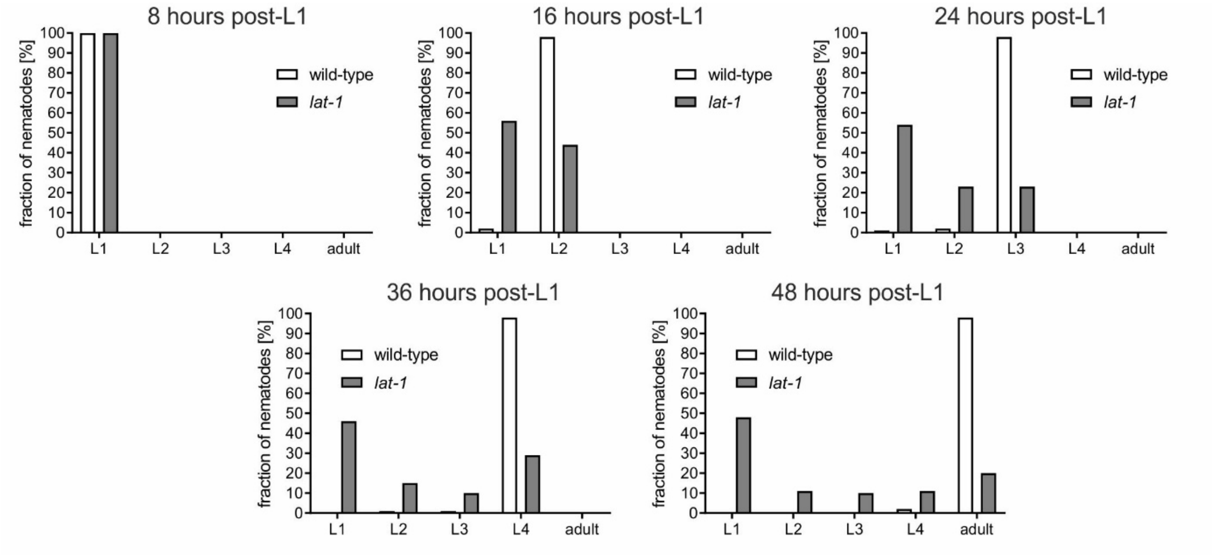


**Figure S1. *lat-1* nematodes display an impaired larval development.** A synchronized L1 population of nematodes was evaluated for progression through the larval stages up to adulthood at 8, 16, 24, 36, and 48 hours post-L1. While wild-type worms show a fast and synchronized progression, some *lat-1* nematodes arrest in early larval stages. However, a small percentage of *lat-1* mutants reach L4/adulthood at the same time as wild-type animals.

| Primer | Sequence 5'-3' |
|---|---|
| lat1_1080F | TTGTTTGTGAGTTTTCGTTTTCGCCTTTATCTTGTTATTTGAATCCGAATCCCTATAGTGAGTCGTATTACATGGTCATA |
| lat1_1081R | ATTGCTGCAGGATCTTTTCTACGGGGTCTGACGCTCAGTG |
| lat1_1082F | GATCCTGCAGCAATGCTTCCGAAAACCTCCGAGTTTTTCG |
| lat1_1083R | GCTACCAGAATCGTTTGGAGCAACGAATAAGTCGTTTTGTTACGTCGCATGCCGTTGAACAGATATAGAAGTTTGTGATT |
| lat1_1084F | CTTAACTGTCAGATTTCAAAAAAAAAAGGA |
| lat1_1085R | CTGTAATTGTCTCATATTTTGATTTTAAAA |
| lat1_1086F | GATCCTGCAGCTTAACTGTCAGATTTCAAAAAA |
| lat1_1087R | GCTACCAGAATCGTTTGGAGCAACGAATAAGTCGTTTTGTTACGTCGCATCTGTAATTGTCTCATATTTTGATTTTAAAAGAGTAAACAC |
| lat1_1088F | TTTGAATCCGAAAAAGTCTCAAAAAGCAAA |
| lat1_1089R | AAAAGGCAAATTTGAAAAGTGTTGTTGGCT |
| lat1_1090F | GATCCTGCAGTTTGAATCCGAAAAAGTCTCAAAAAGCAAA |
| lat1_1091R | GCTACCAGAATCGTTTGGAGCAACGAATAAGTCGTTTTGTTACGTCGCATAAAAGGCAAATTTGAAAAGTGTTGTTGGC |
| lat1_1365F | CGGATGAAAGTGGAACCATCTC |
| lat1_1394F | GCTCCAAACGATTCTGGTAGCTTGTC |
| Oligo d(T)-Anchor Primer | GACCACGCGTATCGATGTCGACTTTTTTTTTTTTTTTTV<br>(V=A, C or G) |
| PCR Anchor Primer | GACCACGCGTATCGATGTCGAC |

**Table S1. Sequences of primers used in the study.**

| GEO Accession | SRA run | raw reads | uniquely aligned raw reads [%] |
|---|---|---|---|
| GSM1862268 | SRR2185654 | 2 x 57999404 | 94.52 |
| GSM1862269 | SRR2185655 | 2 x 40607164 | 94.76 |
| GSM1862270 | SRR2185656 | 2 x 49090847 | 94.62 |

**Table S2. Accession numbers and mapping statistics for transcript analysis of *lat-1* variants.**

| Exon | Chromosome | Start | End |
|---|---|---|---|
| 1 | II | 8805410 | 8805979 |
| 2 | II | 8805628 | 8805979 |
| 3 | II | 8846108 | 8846118 |
| 4 | II | 8890831 | 8890855 |
| 5 | II | 8896750 | 8896987 |
| 6 | II | 8896839 | 8896987 |
| 7 | II | 8896841 | 8896987 |
| 8 | II | 8899797 | 8899984 |
| 9 | II | 8899798 | 8899984 |
| 10 | II | 8903278 | 8903494 |
| 11 | II | 8903388 | 8903494 |
| 12 | II | 8903300 | 8903521 |
| 13 | II | 8903607 | 8903719 |
| 14 | II | 8903832 | 8903929 |
| 15 | II | 8903844 | 8903929 |
| 16 | II | 8903844 | 8903930 |
| 17 | II | 8903913 | 8903929 |
| 18 | II | 8903916 | 8903929 |
| 19 | II | 8904012 | 8904492 |
| 20 | II | 8904012 | 8905264 |
| 21 | II | 8905967 | 8906846 |
| 22 | II | 8906060 | 8906846 |
| 23 | II | 8906575 | 8906846 |
| 24 | II | 8906914 | 8906981 |
| 25 | II | 8906915 | 8907289 |
| 26 | II | 8907152 | 8907287 |
| 27 | II | 8907152 | 8907288 |
| 28 | II | 8907152 | 8907289 |
| 29 | II | 8907725 | 8908630 |
| 30 | II | 8907725 | 8908636 |
| 31 | II | 8964794 | 8964892 |

**Table S3. Exon positions of *lat-1* variants analyzed in this study.** Given are the identified exons for the *lat-1* gene (numbering refers to the reference *C. elegans* genome (WBcel235)).
